# Spatial structure arising from chase-escape interactions with crowding

**DOI:** 10.1101/470799

**Authors:** Anudeep Surendran, Michael J Plank, Matthew J Simpson

## Abstract

Movement of individuals, mediated by localised interactions, plays a key role in numerous processes including cell biology and ecology. In this work, we investigate an individual-based model accounting for various intraspecies and interspecies interactions in a community consisting of two distinct species. In this framework we consider one species to be *chasers* and the other species to be *escapees*, and we focus on chase-escape dynamics where the chasers are biased to move towards the escapees, and the escapees are biased to move away from the chasers. This framework allows us to explore how individual-level directional interactions scale up to influence spatial structure at the macroscale. To focus exclusively on the role of motility and directional bias in determining spatial structure, we consider conservative communities where the number of individuals in each species remains constant. To provide additional information about the individual-based model, we also present a mathematically tractable deterministic approximation based on describing the evolution of the spatial moments. We explore how different features of interactions including interaction strength, spatial extent of interaction, and relative density of species influence the formation of the macroscale spatial patterns.

## Introduction

Movement of, and interaction between individuals is fundamental to various processes in cell biology and ecology^1–3^. These interactions are generated by a number of factors such as the release and detection of chemical signals, mechanical forces, as well as chase and escape dynamics between animals^4–7^, and such interactions can be associated with attractive or repulsive bias^8, 9^. Typically, chase-escape interactions in animal communities involve interspecies interactions between two species where the chasers are biased to move towards the escapees, and the escapees are biased to move away from the chasers^7, 10^. Such chase-escape interactions are relevant to the development of skin patterns in zebrafish involving the movement of two types of pigment cells: melanophores and xanthophores. Here, melanophores tend to move away from xanthophores as xanthophores pursue the melanophores^11^. An example of chase-escape interaction among a single species is in the case of locusts. Cannibalism within the population is an important source of nutrients for locusts. Here, individual locusts tend to escape in response to the threat of being attacked by its conspecifics. Concurrently, they pursue others in the migratory band in search of nutrients^12–14^. Another example of chase-escape dynamics includes herding behaviour, such as in the case of dogs herding sheep^15–17^. Neighbour dependent movement and directional bias can act in unison to generate distinct spatial patterns in the cell population such as cellular aggregation^18^. Similarly, interactions in animal herds or bird flocks can also lead to the creation of spatial patterns^19^. These observations suggest that individual level interactions have significant implications in the macroscale outcomes and motivates us to develop mathematical models to study how the directional movement of individuals as they respond to cues from the neighbourhood, leads to the development of spatial structure.

Mathematical modelling frameworks used to describe the movement of individuals within a population include both discrete individual-based models (IBM) and continuum models that are often described in terms of differential equations^20–22^. To understand the impact of key factors relating to motility, in this work we focus solely on movement processes and neglect proliferation and mortality. Many previous studies focusing on spatial moment dynamics were originally developed to study plant ecology where birth and death are the key mechanisms and motility is irrelevant^23–26^. Recently, this framework has been extended to model cell biology experiments where simultaneous movement and proliferation drives the formation of spatial structure^27, 28^. In our previous work^29^, we considered a general spatial moment model of birth, death and movement process which is relevant to experimental cell biology where motility and proliferation mechanisms act in unison to generate the spatial structure. Development of spatial structure in conservative communities, solely due to motility and interactions such as chase-escape interactions is an interesting problem and to the best of our knowledge, has not been explored in a spatial moment dynamics framework. Therefore, the present work can be viewed as a particular application of our previous work^29^, where we now focus exclusively on the role of motility, and motility bias in a conservative population and we systematically explore how these interactions give rise to spatial structure in the absence of proliferation and death.

In this work we use a lattice-free **IBM** since this approach allows movement in any direction^30^. We also employ interaction kernels to specify the interactions between individuals^31^. Under this model, individuals are either attracted towards or repelled from neighbouring individuals, depending on their species and distance. Unlike models with strict volume exclusion, here there is no absolute restriction on the location an individual occupies and the strength of the interaction is weighted by a distributed interaction kernel. One of the novel features of our study is the extension of previous structure-forming mechanisms of neighbour-dependent directional bias in single species having identical intrinsic properties^9, 27^ to two species with differential attractive/repulsive effects on one another. This significantly expands the types of mechanism that can be modelled, because with a single species it is either attraction or repulsion (or neutral), but with two species there are four types of interaction each of which can be either attraction, repulsion or neutral. In this work the four interactions are: chaser-chaser (c-c); chaser-escapee (c-e); escapee-chaser (e-c); and escapee-escapee (e-e).

Spatial moment analysis is a continuum approximation that tracks the dynamics of the density of individuals, pairs, triples and so on, to describe the spatial correlation present in a population^24, 32, 33^. Classical macroscale continuum models such as the logistic growth model, disregard the influence of spatial structure by invoking a mean-field approximation such that the interactions between individuals are in proportion to their average density^34^. Spatial moment analysis is advantageous over traditional mean-field analysis because it accounts for short-range, neighbour-dependent interactions and spatial correlations, thereby allowing for the analysis of clustered and segregated populations^31^.

### Stochastic, individual-based model

We consider a community of two species: *chasers* and *escapees*. We construct the IBM for a spatially homogeneous environment, where the probability of finding an individual in a small region is independent of the location of that region. Hence, the IBM is relevant for communities that do not involve macroscopic density gradients meaning that we do not consider moving fronts or travelling waves^35, 36^. We consider a community with *N*_*c*_ chasers and *N*_*e*_ escapees, distributed over a continuous two-dimensional finite space, Ω ∈ ℝ^2^. At any instant of time *t*, the state of the IBM is characterised by the locations of the two types of individuals **x**_*n*_ ∈ Ω, where *n* = 1, 2, …, *N*_*c*_ + *N*_*e*_. The intrinsic movement rates of chasers and escapees are *m*_*c*_ and *m*_*e*_, respectively. We consider a continuous time Markov process where the probabilities that an isolated chaser or an isolated escapee moves during a short time interval of duration *δt* is *m*_*c*_*δt* and *m*_*e*_*δt*, respectively.

An important feature of the IBM is the neighbour dependent directional bias where movement directions of individuals are influenced by the interactions with other individuals in the neighbourhood. The interaction between two chasers at a displacement *ξ* is specified by the bias interaction kernel, *ω*_*cc*_(|*ξ*|). We choose these kernels to be two-dimensional Gaussian functions, given by,

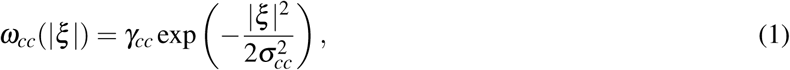

where *γ*_*cc*_ and *σ*_*cc*_ *>* 0 represent the interaction strength and the spatial extent of interaction, respectively. The intraspecies interaction of escapees as well as interspecies interactions between chasers and escapees are specified by similar interaction kernels, *ω*_*ee*_(|*ξ*|), *ω*_*ce*_(|*ξ*|), and *ω*_*ec*_(|*ξ*|), respectively. The Gaussian shape of the bias kernel means that individuals interact strongly with their immediate neighbours while interactions between more distant pairs are negligible. We ensure that the size of the computational domain is sufficiently large relative to *σ*_*ij*_ so that pairs of agents only interact in some local neighbourhood of the domain. The interaction kernel in Equation 1 is symmetric, implying that we obtain steady spatial patterns. However, our modelling framework is sufficiently general to incorporate asymmetric kernels if that were of interest. In this case, asymmetric kernels could lead to other types of spatial structures such as stable moving configurations of chasers following escapees^14^.

The sign of the interaction strength determines the nature of the interaction. A negative bias strength corresponds to attractive interactions while positive bias strength corresponds to repulsive interactions, and we refer to these interactions are as *chase* and *escape* interactions, respectively. The relationship between the directional bias and the nature of the local interactions is best illustrated by considering a few of the possible scenarios in a community of chasers and escapees as shown in Figure 1. In Figure 1(a), we see that a reference chaser does not have a preferred direction since they feel no bias from neighbouring chasers, because *γ*_*cc*_ = 0. For the intraspecies interactions of escapees, shown in Figure 1(b), a positive *γ*_*ee*_ results in the reference escapee moves away from other escapees. Similarly, for the interspecies interaction of chasers and escapees, shown in Figure 1(c)-(d), a negative *γ*_*ce*_ results in chaser attracted to escapees, whereas a positive *γ*_*ec*_ results in escapees repelled from chasers. In this way, we have the flexibility of choosing the interspecies interaction to be asymmetric and later we will show that this flexibility can give rise to some very interesting results.

**Figure 1.**
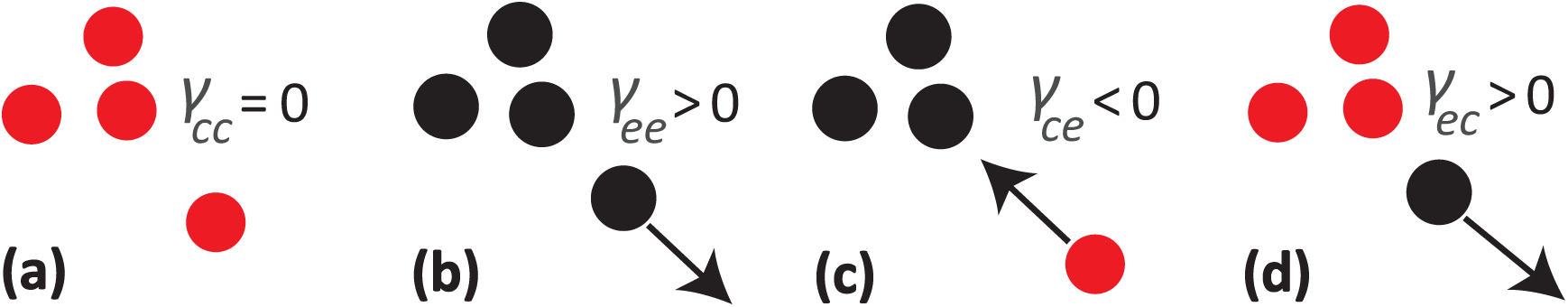
Schematic representation of some of the possible interactions arising in a community of chasers (red) and escapees (black). The bias strength of intraspecies interactions of chasers and escapees are *γ*_*cc*_ and *γ*_*ee*_, respectively. The bias strengths of interspecies interactions are *γ*_*ce*_ and *γ*_*ec*_, respectively. **a** No intraspecies interaction between chasers. **b** Intraspecies repulsion between escapees. **c** Chasers attracted to escapees. **d** Escapees repulsed from chasers.

We define a function, *Q*(**x**, *t*), called the *bias landscape*, which allows us to quantity crowdedness and interactions at location **x** and time *t*. The bias landscape for a chaser is defined as a sum of bias kernels around each individual in the community:

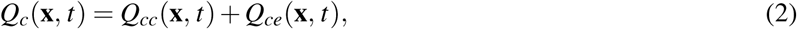

where 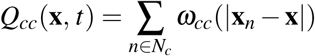 and 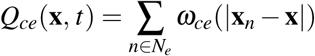. The bias landscape for escapees is defined very similarly *Q*_*e*_(**x**, *t*) = *Q*_*ec*_(**x**, *t*) + *Q*_*ee*_(**x**,*t*). For any individual *n*, we define the net bias vector to be

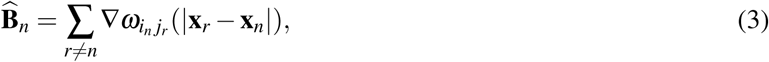

where *i, j* ∈ {*c, e*}. Two key features of the bias vector are the magnitude, 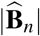, and angular direction, arg 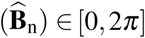. The preferred direction of movement is taken to be the angular direction of the net bias vector and this allows us to bias the direction of movement of individuals in response to the local crowdedness. The definition of the bias vector, Equation 3, shows that the bias vector is the negative gradient of the bias landscape meaning that each individual is biased to move in the direction of the steepest descent on the surface. The strength of the bias is determined by the steepness of the bias landscape at any particular individual’s location.

The construction of the bias landscape and bias vectors in a community of chasers and escapees is shown in Figure 2. The level curves of the intraspecific components of the bias landscapes, *Q*_*cc*_(**x**, *t*) and *Q*_*ee*_(**x**, *t*), are shown in Figure 2(a)-(b), respectively. The individual bias vectors point in the direction of steepest descent. In this example we assume no interspecies interaction of chasers (*γ*_*cc*_ = 0), meaning that their bias vector is 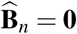. This explains why there are no vectors in Figure 2(a), and so the direction of movement of chasers is unaffected by other chasers. In contrast, the bias vectors for escapees in Figure 2(b) are directed away from each other due to their intraspecies repulsion. The net bias landscape for chasers, *Q*_*c*_(**x**, *t*), shown in Figure 2(c), includes the additional interspecies contribution from escapees so that the chasers are attracted to escapees. Note that, the chaser at approximately (*x, y*) = (−0.5, −5) is not influenced by the interspecies interaction because it is too far away from the clusters of escapees. Similarly, including the interspecific component of the bias landscape for escapees, shown in Figure 2(d), reorients the escapees’ bias vectors away from any nearby chasers.

**Figure 2.**
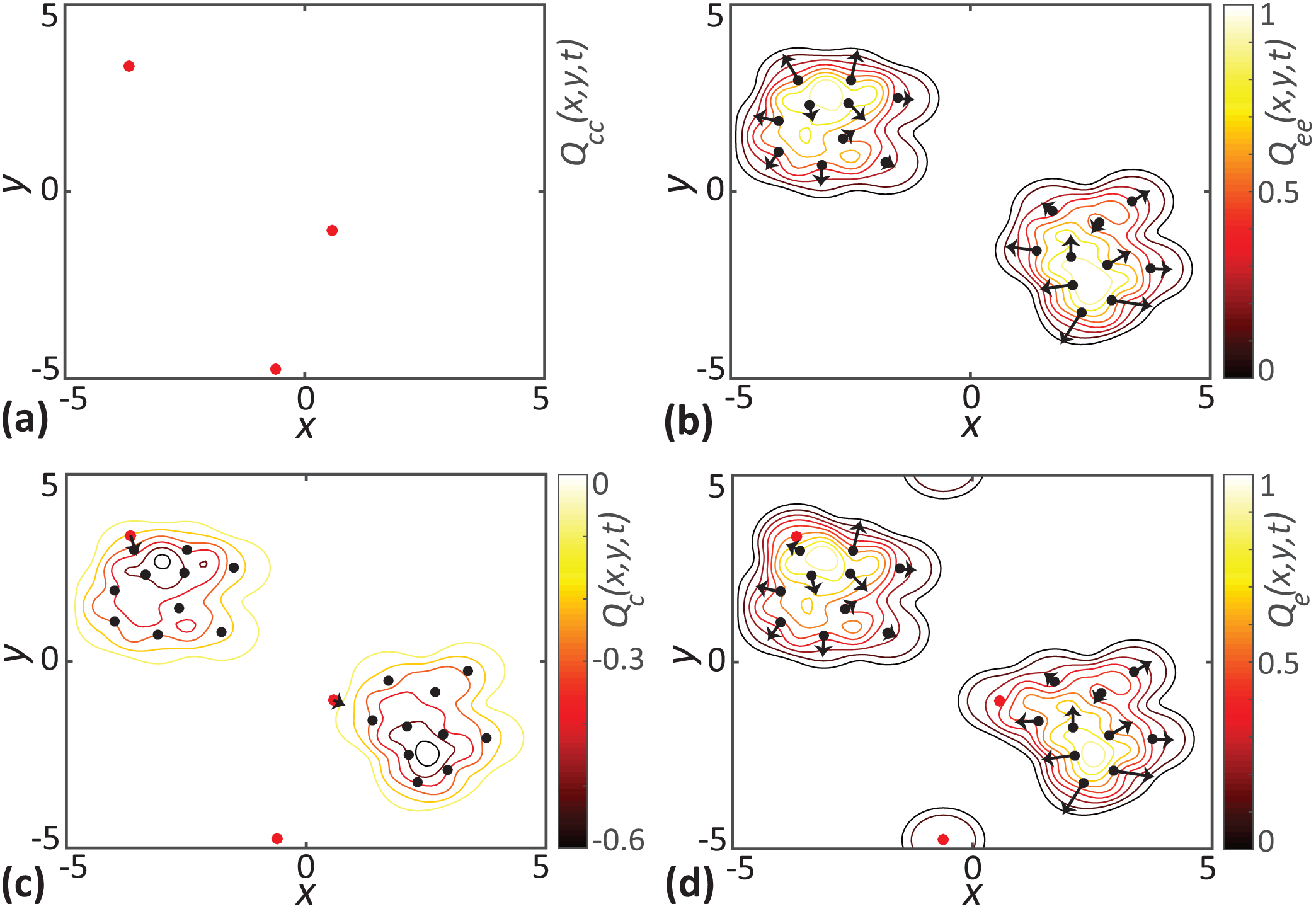
Visualisation of directional bias for a community of chasers (red dots) and escapees (black dots). In each subfigure, **a-d**, the locations of individuals are superimposed with the level curves of the bias landscapes. The bias vectors shown with black arrows represent the preferred direction of individuals. The length of arrows indicate the strength of bias. Results in **a** correspond to intraspecies interactions of chasers only. Results in **b** show intraspecies interactions of escapees only. Results in **c** and **d** show the net interaction for both species. Parameter values are *γ*_*cc*_ = 0.0, *γ*_*ce*_ = −0.2, *γ*_*ec*_ = *γ*_*ee*_ = 0.3 and *σ*_*cc*_ = *σ*_*ce*_ = *σ*_*ec*_ = *σ*_*ee*_ = 0.5. Note that *γ*_*cc*_ = 0, meaning that there are no interactions between chasers in (a), which explains why there is no colour bar on this subfigure.

We now specify the probability density function (PDF) for the displacement that a chaser traverses during a movement event as,

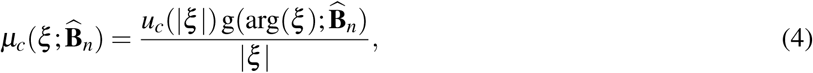

where *u*_*c*_(|*ξ*|) and 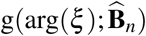 are two independent PDFs. A derivation of the movement displacement PDF is provided in Section 1 of the Supplementary Material. A similar definition holds for the movement displacement PDF of escapees, 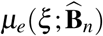. Here, *u*_*c*_(|*ξ*|) is the PDF for the movement distance, |*ξ*| moved by a chaser. For simplicity, we treat *u*_*c*_(|*ξ*|) as a PDF that is independent of the local environment. It is possible to use a constant, deterministic step length in the IBM^28^. However, since we are interested in deriving a continuous mathematical description of the IBM we use a continuous distribution to describe the step length. Note that the expression for 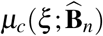 involves a factor of 1*/*|*ξ*| and so evaluating 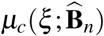 at *ξ* = 0 results in the PDF becoming infinite. To avoid this, we use a non-zero mean 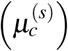 normal distribution with a relatively narrow standard deviation 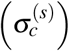, where 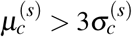 for the movement distance PDF, *u*_*c*_(|*ξ* |). When we evaluate the movement displacement PDF, we set *u*_*c*_(|*ξ* |) = 0 at distances greater than 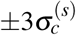 from the mean distance. It is also possible to use other strictly positive distributions for the step length, such as a gamma distribution. However, we choose the truncated Gaussian PDF to be consistent with the previous studies in this field^27, 29^. Provided the PDF has a relatively narrow distribution around the mean and has zero density at the origin, the exact choice of step length distribution does not qualitatively change the results. Now, 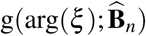 is the PDF describing the angular direction of movement, arg(*ξ*) ∈ [0, 2*π*], of a chaser due to the presence of other individuals in the neighbourhood. We choose the PDF for the angular direction of movement to be a von Mises distribution, with mean direction 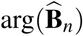 and concentration parameter given by 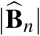,

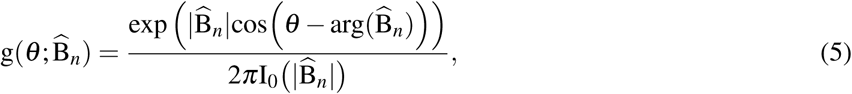

where *I*_0_ is the zero order modified Bessel function. In the absence of net bias,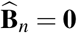, the von Mises distribution reduces to the uniform distribution so that there is no preferred direction of movement^27^. This choice of PDF means that the likelihood of individuals moving in the preferred direction, 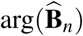, increases with the magnitude of the bias vector. Individuals with large 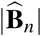, are more likely to move in the direction of 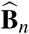. The direction of movement of individuals with a weaker bias becomes almost uniformly distributed. Note that, the magnitude of 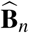 does not affect the distance moved.

We implement the IBM using the Gillespie algorithm^37^ with an initial condition where individuals of each species are distributed uniformly at random in the domain. An alternative simulation method would be to use a constant time-stepping approach, but this can be less accurate because there is a possibility that multiple events occur in the same time step^38^. Note that, our model considers only the movement direction to be neighbour dependent, and movement rates are chosen to be independent of neighbourhood interactions. In general, our framework could be extended to allow both the movement rates and movement direction to depend upon the crowding surface. However, we choose to work with the simplest implementation where the movement rate does not depend upon the crowding surface since this simpler mechanism gives rise to very interesting spatial patterns.

### Numerical simulation of the individual-based model

In our IBM simulations, we consider a community of chasers and escapees with population sizes *N*_*c*_ and *N*_*e*_, respectively, distributed uniformly at random in a domain of size *L*× *L*. The intrinsic movement rates of chasers and escapees are *m*_*c*_ and *m*_*e*_, respectively. The net movement rate of the community is given by,

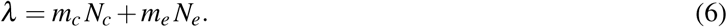

The time interval between successive movement events is exponentially distributed with mean 1*/λ*. When a motility event takes place, an individual is chosen to move at random. Depending upon whether the chosen individual is a chaser or escapee, it moves a displacement *ξ* specified by either of the movement displacement PDFs, 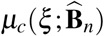, and 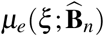. We employ periodic boundary conditions over the *L*× *L* domain in all of our simulations. Typically, the length scales over which interactions take place between individuals are much smaller compared to the overall domain size (*σ*_*ij*_ ≪ *L*). Under these circumstances, the edge effects are negligible, and the periodic boundary condition is a reasonable choice. Also, many previous works that focus on modelling the motility and interactions in ecological and biological systems employ periodic boundary conditions^9, 14, 16, 29^.

We use spatial moments to characterise the spatial structure in the population. The first spatial moment of chasers is *Z*_1,*c*_ = *N*_*c*_*/L*^2^ and that of escapees is *Z*_1,*c*_ = *N*_*e*_*/L*^2^. We analyse the spatial structure of the community using second spatial moments expressed as auto and cross-PCFs. To compute the auto-PCF of chasers, we choose a reference chaser located at **x**_*n*_ and compute all distances, *r* = |**x**_*r*_ − **x**_*n*_| from all other chasers. We repeat this procedure until each of the remaining *N*_*c*_ − 1 chasers act as a reference individual. The PCF is constructed by counting all the distances that fall into the interval [*r* −*δr/*2, *r* + *δr/*2]. Normalising the bin count by a factor of *N*_*c*_ (*N*_*c*_ − 1)(2*πrδr*)*/L*^2^ gives *C*_*cc*_ such that *C*_*cc*_(*r*) = 1 corresponds to a complete absence of spatial structure among chasers. To compute *C*_*ee*_ and *C*_*ce*_, we repeat this procedure by considering pairs of escapees and pairs involving both chasers and escapees, respectively.

### Spatial moment dynamics

In this section, we derive a macroscale continuum approximation of the IBM by considering the dynamics of spatial moments. For convenience, we present the spatial moment dynamics for a multispecies framework and specific details corresponding to the two species case are presented in Section 2 of the Supplementary Material. The spatial point process defined by the IBM is stationary, meaning that statistics of the population in any observation window are independent of the window’s location^31^. This means that the first spatial moment *Z*_1,*i*_, the average density of species *i*, is independent of location. Since we do not consider the proliferation and death of individuals, the first moment remains constant in time. The second spatial moment, *Z*_2,*ij*_(*ξ, t*), is the average density of pairs composed of individuals of species *i* and species *j*, separated by a displacement *ξ*. The third spatial moment, *Z*_3,*ijk*_(*ξ, ξ′,t*), is defined as the average density of triplets consisting of individuals of species *i*, species *j* and species *k* where the individuals of species *j* and *k* are displaced by *ξ* and *ξ′*, respectively from the individual of species *i*. Note that since the subpopulations are conservative we refer to the average density as *Z*_1,*i*_ whereas the second and third moments are time dependent, so we explicitly show the dependence on distance and time by writing *Z*_2,*ij*_(*ξ, t*) and *Z*_3,*ijk*_(*ξ, ξ′,t*), respectively.

### General spatial moment formulation

To derive the equations governing the dynamics of the second spatial moment, we need to derive the expected net bias vector for an individual of species *i* that is in pair with an individual of species *j*. This depends on the contribution from another individual of species *k* at a displacement *ξ′*. Conditional on a pair consisting of individuals of species *i* and *j* existing at a displacement *ξ*, the probability of an individual of species *k* being present at a displacement *ξ*′ is *Z*_3,*ijk*_(*ξ, ξ* ′,*t*)*/Z*_2,*ij*_(*ξ, t*). The expected net bias vector of an individual of species *i*, conditional on the presence of an individual of species *j* can be computed by multiplying the gradient of bias kernel with the conditional probability and integrating over all possible displacements, giving,

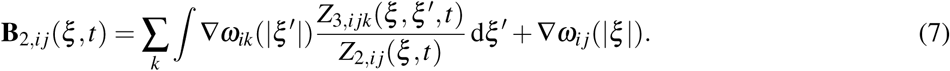

The second term in Equation 7, ∇*ω*_*ij*_(|*ξ* |), accounts for the direct contribution of the individual of species *j* at displacement *ξ*. Now we can write the PDF for the movement of an individual of species *i* over a displacement *ξ′*, conditional on the presence of an individual of species *j* at a displacement *ξ* as,

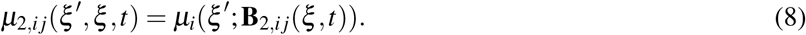

We now consider the dynamics of the second spatial moment by considering the change in the density of pairs of individuals at a displacement *ξ* as a result of the movement of individuals. An existing pair of individuals at displacement *ξ* can be lost if one of the individuals of the pair moves. Similarly, a new pair with displacement *ξ* is formed when one of the individuals from an existing pair at a displacement *ξ* + *ξ′* moves a displacement *ξ′*. Accounting for these possibilities means that the time evolution of the second moment is given by,

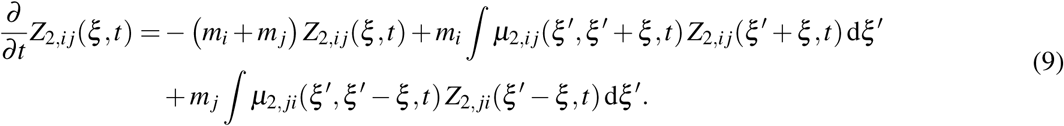

Given estimates of the first and second moments, we can describe the spatial structure of the community in terms of a pair correlation function (PCF), *C*_*i j*_(*r,t*), expressed as a function of separation distance, *r* = |*ξ* |, given by,

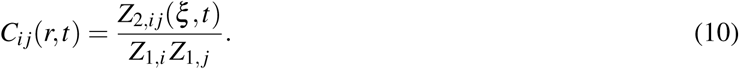

The normalisation of the second spatial moment by a factor of *Z*_1,*i*_ *Z*_1, *j*_ in Equation 10 ensures that *C*_*ij*_(*r,t*) = 1 in the absence of spatial structure. If *C*_*ij*_(*r,t*) *>* 1, we have more pairs of an individual of species *i* separated by a distance *r* from an individual of species *j* than if they were in a spatially random configuration. This is referred to as a clustered spatial structure. If *C*_*ij*_(*r,t*) *<* 1, we have fewer pairs of individuals at a separation distance *r* than if they were in a spatially random configuration and this is known as a regular spatial pattern. The PCF is called an auto-PCF when describing the density of pairs involving individuals of the same species and cross-PCF when individuals are from two different species^39^.

The dynamics of the second spatial moment depend on the third spatial moment via Equation 7. To obtain a closed system of equations, we use a moment closure scheme to approximate the third moment as a function of first and second moments^40, 41^. Different closure schemes have been used in the literature including the power-1 closure, power-2 closure and the power-3 Kirkwood superposition approximation^42^. A comparison of the performance of various closure methods for the kinds of problems we consider is summarised in Section 3 of the Supplementary Material. We find that the asymmetric power-2 closure^26^ given by,

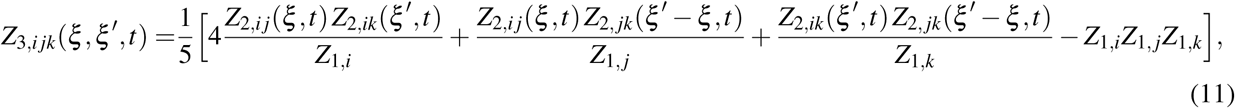

provides the best results for the range of parameters we consider.

### Summary statistics

In our analysis of both the IBM and spatial moment model, we focus on examining sufficienty long-time simulations so that we can treat the higher moments and *C*_*ij*_(*r,t*) as quasi-steady quantities^43^. Under these circumstances, we specify *C*_*ij*_(*r*) as a function of *r* only. To summarise the key features of the various possible combinations of spatial structure, we need a simplified measure of spatial structure. Plotting and comparing the various PCFs as a function of distance for each parameter value becomes increasingly challenging as we consider a large parameter space. Under these circumstances, it is more convenient to use a simple measure of spatial structure that expresses the type and extent of the spatial structure as a scalar quantity. A summary statistic based on the difference in the area under the curve of the PCF and the area under a curve made by a constant pair correlation function for a randomly distributed population, *C*_*ij*_(*r*) = 1, is one convenient way to simply express the nature and extent of the spatial structure as a single number. In this work we use,

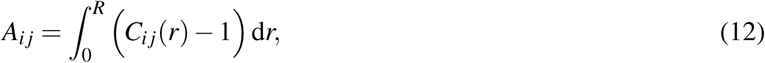

as such a summary statistic where *A*_*ij*_ quantifies the spatial structure between of species *i* and *j* as a function of *C*_*ij*_(*r*). Binny et al. (2016)^9^ use a similar measure of spatial structure to describe spatial patterns in a single species framework. Here we use a slightly more general summary statistic since we are interested in populations composed of multiple species. Our definition of *A*_*ij*_ is such that, if individuals of species *i* are more (less) likely to be located in proximity to individuals of species *j*, then *A*_*ij*_ *>* 0 (*A*_*ij*_ *<* 0). If there is no correlation between locations of individuals of species *i* and *j* then *A*_*ij*_ = 0. Even though, it is possible to have a positive correlation (*C*_*ij*_(*r*) *>* 1) at some scales and negative correlation (*C*_*ij*_(*r*) *<* 1) at other scales, in practice we do not observe this situation. We find that *A*_*ij*_ is insensitive to *R* when we choose *R* to be sufficiently large since the PCFs approach unity for large separation distances.

### Numerical methods for spatial moment dynamics

The numerical solution for the dynamical equation of the second spatial moment, Equation 9, is computed using the forward Euler method. This scheme involves a spatial discretisation of the displacement, *ξ* = (*ξ*_*x*_, *ξ*_*y*_), using a square grid with constant spacing, Δ*ξ*, over the domain {−*ξ*_max_ *≤ ξ*_*x*_, *ξ*_*y*_ *≤ ξ*_max_}. Here, *ξ*_max_ is chosen to be sufficiently large such that the second spatial moments are given by a mean field condition at the domain boundaries, giving, *Z*_2,*ij*_(*ξ, t*) ≈ *Z*_1,*i*_ × *Z*_1,*j*_. The integral terms in Equation 7 and Equation 9 are approximated using the trapezoid rule. To compute these integrals, we need to evaluate *Z*_2,*ij*_(*ξ ±ξ* ′,*t*) for different combinations of *ξ* and *ξ* ′. Some of these combinations require values of *Z*_2,*ij*_(*ξ ± ξ* ′,*t*) that lie beyond the computational domain. Whenever this scenario arises, we approximate those terms with the values of *Z*_2,*ij*_(*ξ, t*) at the boundary, *Z*_2,*ij*_((*ξ*_max_, *ξ*_max_),*t*).

The initial condition used for solving the spatial moment dynamics is *Z*_2,*ij*_(*ξ*, 0) = *Z*_1,*i*_ × *Z*_1,*j*_. We use a constant grid spacing Δ*ξ* = 0.2, time step d*t* = 0.1 and *ξ*_max_ = 8 in all our results. Additional results (not shown) confirms that this choice of spatial and temporal discretisation is sufficient to produce grid-independent results. We compare the numerical solution of the spatial moment dynamics model with the IBM simulation results by computing the auto-PCF and cross-PCF 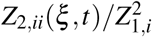 and *Z*_2,*i j*_(*ξ, t*)*/*(*Z*_1,*i*_ *Z*_1,*j*_), respectively.

## Results

Here, we explore how different features of the intraspecies and interspecies interactions influence the spatial structure of a two species community using the IBM and spatial moment models. We examine the influence of variation in factors such as interaction strength, spatial extent of interaction and relative density of species, in determining the macroscale spatial patterns. Our results show that, at a sufficiently large time, *t* = 20, a steady spatial pattern establishes in the community. The evolution of the spatial structure of the community to steady state is shown in Section 4 of the Supplementary Material. Since our interest lies in understanding the long-time impact of individual-level interactions, we focus on presenting and interpreting steady-state solutions of our models. While the time required to attain a steady-state depends on the specific initial distribution of individuals in the domain, the actual steady-state spatial pattern formed is independent of the initial configuration. To illustrate, we provide additional results in Section 5 of the Supplementary Material. In all our simulations, the individuals are initially distributed uniformly at random. This choice helps in reducing the computational time and comparing the influence of different types of interactions. A summary of the key parameters in the simulations is given in Table 1. Before presenting the details about how different features of interactions influence the spatial structure, we illustrate the types of spatial structure that could arise in a community of chasers and escapes by showing the snapshots of locations of individuals from IBM in Figure 3. A community with a complete absence of spatial structure is shown in Figure 3(a), where there is no correlation between the locations of chasers and escapees. Figure 3(b) shows the interspecies clustering of chasers and escapees, formed due to the interspecies attraction of chasers towards escapees. Here we see more chasers and escapees in close vicinity of each other than in a spatially random configuration. Figure 3(c) shows the interspecies regular pattern formed due to interspecies chase-escape interactions. In this case, the repulsive bias of escapees from chasers is stronger than the attraction of chasers to escapees. Hence we see that escapees are placed away from chasers. Note that, the identification of the spatial structure of a community purely through visual inspection can be challenging in certain cases. Under these scenarios, PCFs are very useful in revealing the type of spatial structure and quantifying it.

**Table 1.**
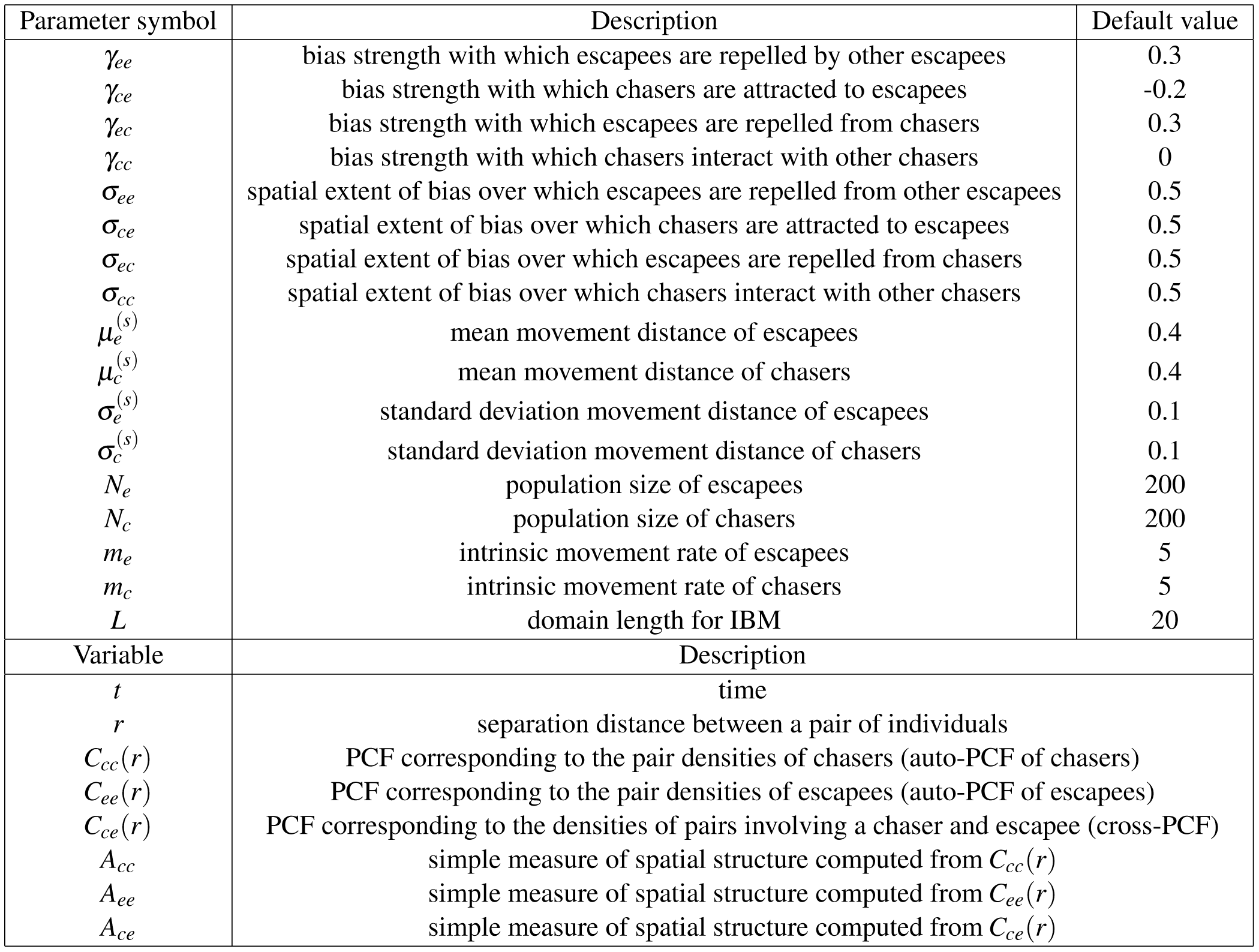
Description of model parameters and variables.

**Figure 3.**
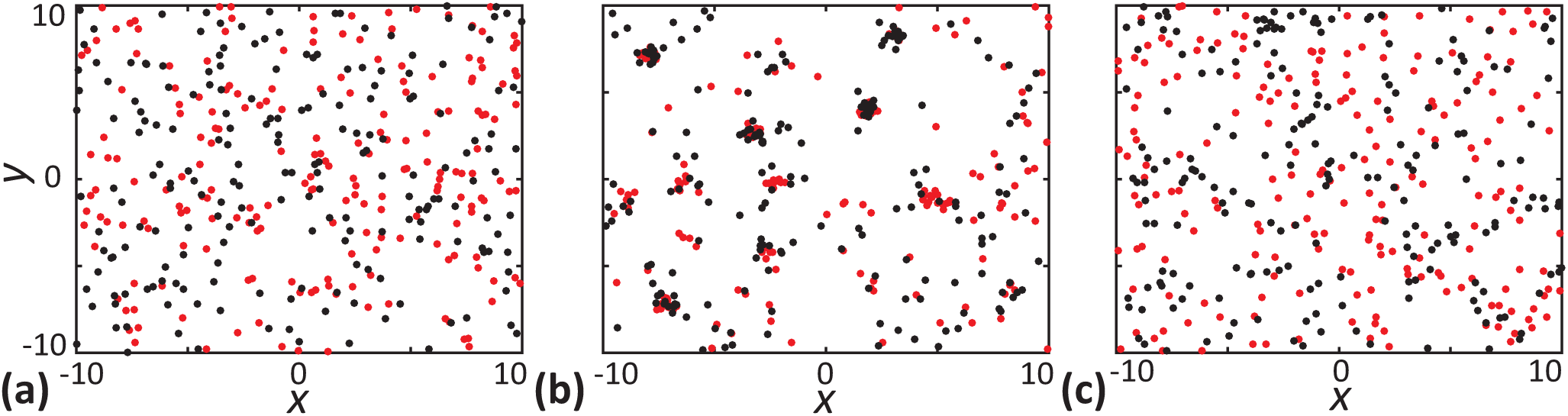
Snapshots of the locations of chasers and escapees to illustrate the types of spatial structure arise in our model. **a** Complete absence of spatial structure (*γ*_*ce*_ = *γ*_*ec*_ = 0). **b** Interspecies cluster spatial structure (chasers and escapees form aggregates, *γ*_*ce*_ = −0.25, *γ*_*ec*_ = 0, *γ*_*ee*_ = −0.06). **c** Interspecies regular pattern (chasers and escapees are segregated, *γ*_*ce*_ = −0.1, *γ*_*ec*_ = 0.3). Other parameter values are *γ*_*cc*_ = *γ*_*ee*_ = 0, *σ*_*cc*_ = *σ*_*ce*_ = *σ*_*ec*_ = *σ*_*ee*_ = 0.5, *m* = *m* = 5.0, 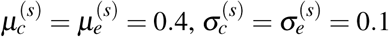.

### Effect of varying the interaction strength

Here, we investigate how variations in bias interaction strength impact the spatial structure. In our simulations, we consider a two species community of chasers and escapees with population sizes, *N*_*c*_ = 200 and *N*_*e*_ = 200, distributed according to a spatial Poisson process. We systematically vary the interaction strengths *γ*_*ce*_, *γ*_*ec*_, and *γ*_*ee*_ to explore the role of different interactions. For simplicity, we assume *γ*_*cc*_ = 0, which means that the chasers have no intraspecific interactions. We compute the PCFs, *C*_*cc*_(*r*), *C*_*ce*_(*r*), and *C*_*ee*_(*r*) along with simple measures of spatial structure, *A*_*cc*_, *A*_*ce*_ and *A*_*ee*_, to quantify the resulting spatial patterns generated in the community, as shown in Figure 4.

**Figure 4.**
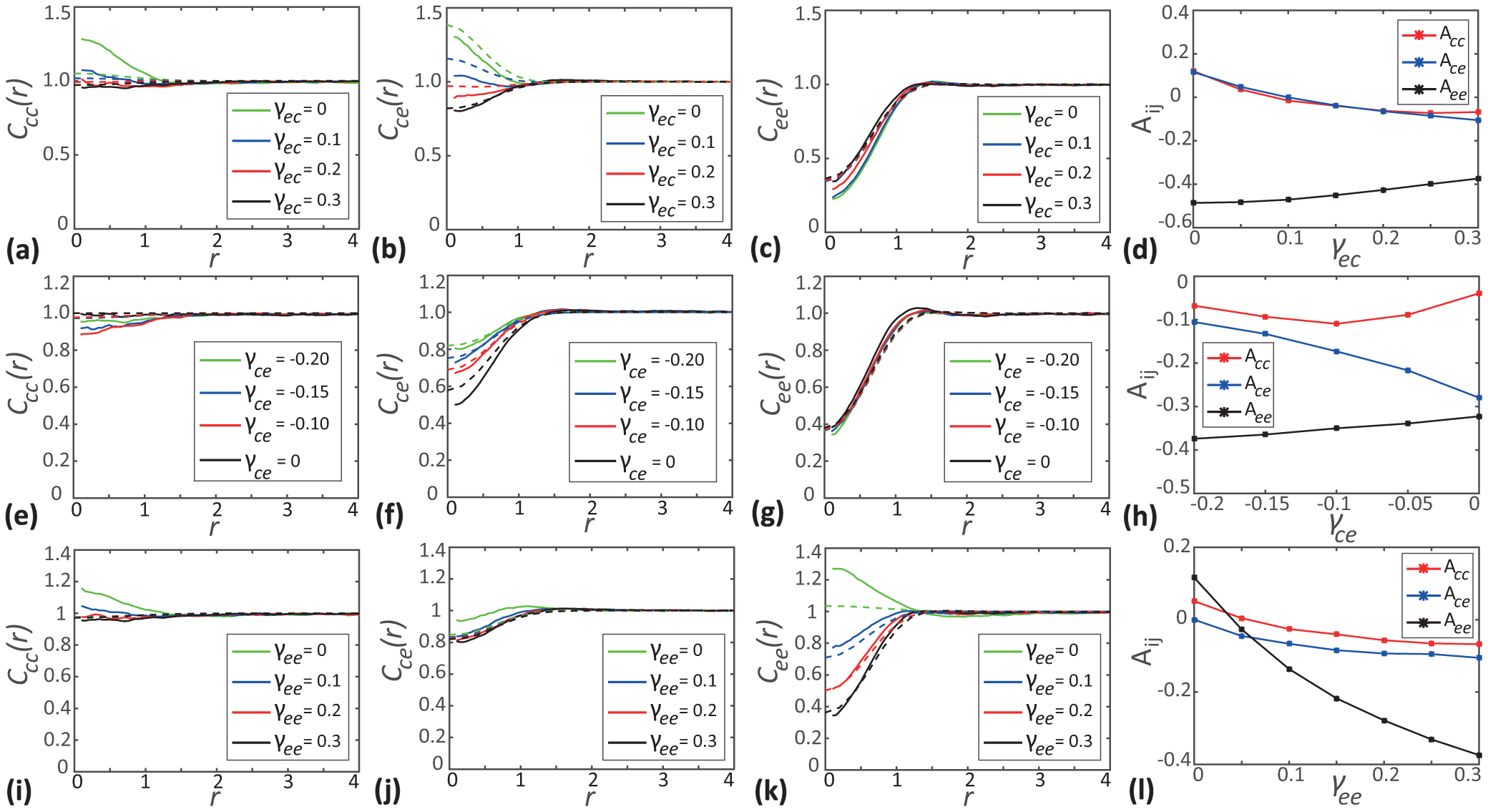
Effects of varying the interaction strength. **a-c** show the PCFs *C*_*cc*_(*r*), *C*_*ce*_(*r*) and *C*_*ee*_(*r*) as a function of separation distance for different choices of *γ*_*ec*_. **d** show simple measures of spatial structure *A*_*cc*_, *A*_*ce*_ and *A*_*ee*_ as a function of *γ*_*ec*_. **e-g** shows *C*_*i j*_(*r*) for different choices of *γ*_*ce*_. **h** shows *A*_*i j*_ as a function of *γ*_*ce*_. **i-k** show *C*_*i j*_(*r*) for different choices of *γ*_*ee*_. **l** shows *A*_*i j*_ as a function of *γ*_*ee*_. In the subfigures showing the PCFs as a function of *r*, solid curves show the averaged results from 1000 identically prepared realisations of the IBM, whereas dashed curves correspond to results from spatial moment dynamics model. In all cases *C*_*i j*_(*r*) and *A*_*i j*_ are evaluated at *t* = 20. Parameter values are *γ*_*cc*_ = 0, *σ*_*cc*_ = *σ*_*ce*_ = *σ*_*ec*_ = *σ*_*ee*_ = 0.5, *m*_*c*_ = *m*_*e*_ = 5.0, 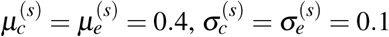.

In Figure 4(a)-(d) we vary the bias interaction strength, *γ*_*ec*_, which controls the strength with which escapees are repelled from chasers. When escapees are not repelled from chasers (*γ*_*ec*_ = 0), the chasers move close to escapees due to their attractive bias, leading to the accumulation of chasers around escapees. This results in intraspecies clustering of chasers (*C*_*cc*_(*r*) *>* 1) as well as interspecies cluster formation (*C*_*ce*_(*r*) *>* 1). This is also confirmed by, simple measures of spatial structure, *A*_*cc*_ and *A*_*ce*_ being positive when *γ*_*ec*_ = 0. We also see a regular pattern among escapees due to their intraspecies repulsion (*C*_*ee*_(*r*) *<* 1 and *A*_*ee*_ *<* 0). When escapees are repelled from chasers (*γ*_*ec*_ *>* 0), the clustering we previously observed weakens as the repulsion counteracts the attractive bias of chasers. For a sufficiently large choice of *γ*_*ec*_, here *γ*_*ec*_ = 0.3, spatial structure changes becomes regular due to the overall segregation of chasers and escapees (*C*_*cc*_(*r*) *<* 1 and *A*_*cc*_ *<* 0 as well as *C*_*ce*_(*r*) *<* 1 and *A*_*ce*_ *<* 0). In spite of the intraspecies repulsion between escapees, the enhanced interspecies repulsion from chasers, force the escapees to come closer to each other. This results in weakening of the regular spatial structure of escapees as *γ*_*ec*_ increases.

In Figure 4(e)-(h) we vary *γ*_*ce*_, which controls the strength with which chasers are attracted to escapees. When chasers are not attracted to escapees (*γ*_*ce*_ = 0), the movement of chasers is unbiased so their distribution remains spatially random (*C*_*cc*_(*r*) = 1). In this case, there is no effect opposing the repulsion of escapees from chasers. This results in escapees staying away from chasers, forming an interspecies regular pattern, as indicated by *C*_*ce*_(*r*) *<* 1 for small distances and *A*_*ce*_ *<* 0. Also, intraspecies repulsion among escapees contributes to the regular pattern of escapees, indicated by *C*_*ee*_(*r*) *<* 1 and *A*_*ee*_ *<* 0. When chasers are attracted to escapees (*γ*_*ce*_ *<* 0), an intraspecies regular spatial structure emerges among chasers (*C*_*cc*_(*r*) *<* 1 and *A*_*cc*_ *<* 0). This is because chasers tend to move towards escapees and the regular arrangement of escapees forces chasers to occupy spaces around escapees. As the attraction strength is increased (*γ*_*ce*_ more negative), the interspecies regular pattern becomes less pronounced, since the enhanced attractive bias starts to counteract the repulsion from escapees. As *γ*_*ce*_ is made more negative, *A*_*ce*_ gets closer to zero also confirm this behaviour.

In Figure 4(i)-(l) we vary *γ*_*ee*_, which controls the strength with which escapees are repelled by other escapees. As we increase the intraspecies repulsion strength of escapees, we observe a strong regular spatial pattern emerges among escapees, as indicated by *C*_*ee*_(*r*) diverges form unity towards zero. The *A*_*ee*_ becomes more negative with the increase in *γ*_*ee*_ also confirms this behaviour. The development of more pronounced intraspecies regular spatial structure among escapees can be attributed to the increased repulsion between escapees as *γ*_*ee*_ increases. Finally, we observe that the accuracy of the spatial moment prediction increases with the increase in the magnitude of the bias strength.

### Effect of varying the spatial extent of interaction

Here, we investigate how varying the range of interactions impacts the formation of spatial structure. The spatial extent of interaction, *σ*_*i j*_ determines the range over which the interactions between individuals are significant. A small choice of *σ*_*i j*_ leads to short-range interactions, whereas a large *σ*_*i j*_ corresponds to long-range interactions between individuals. Again, we consider a two species community of chasers and escapees with *N*_*c*_ = 200 and *N*_*e*_ = 200, respectively. As before we set *γ*_*cc*_ = 0, meaning the interactions in between chasers are switched off.

In Figure 5(a)-(d) we vary the spatial extent of interaction *σ*_*ec*_, which controls the spatial extent over which escapees are repelled from chasers. For small values of *σ*_*ec*_, we observe a small scale intraspecies clustering among chasers as indicated by the PCF *C*_*cc*_(*r*) *>* 1 and the simple measure of spatial structure *A*_*cc*_ *>* 0. We also observe interspecies clustering between chasers and escapees (*C*_*ce*_(*r*) *>* 1) as well as a regular spatial pattern between escapees (*C*_*ee*_(*r*) *<* 1) at small values of *σ*_*ec*_. Our results showing that *A*_*ce*_ *>* 0 and *A*_*ee*_ *<* 0 also confirm this behaviour for small *σ*_*ec*_. As *σ*_*ec*_ increases, the intraspecies clustering of chasers and interspecies clustering becomes weaker and changes to a regular spatial structure at a sufficiently large values of *σ*_*ec*_. As the range over which escapees are repelled increases, the escapees experience stronger repulsion due to the influence of more distant chasers resulting in the segregation of chasers and escapees. We observe that both *A*_*cc*_ and *A*_*ce*_ change sign from a positive value at *σ*_*ec*_ = 0.25 to negative values as we increase *σ*_*ec*_ confirms this trend of development of regular spatial structure.

**Figure 5.**
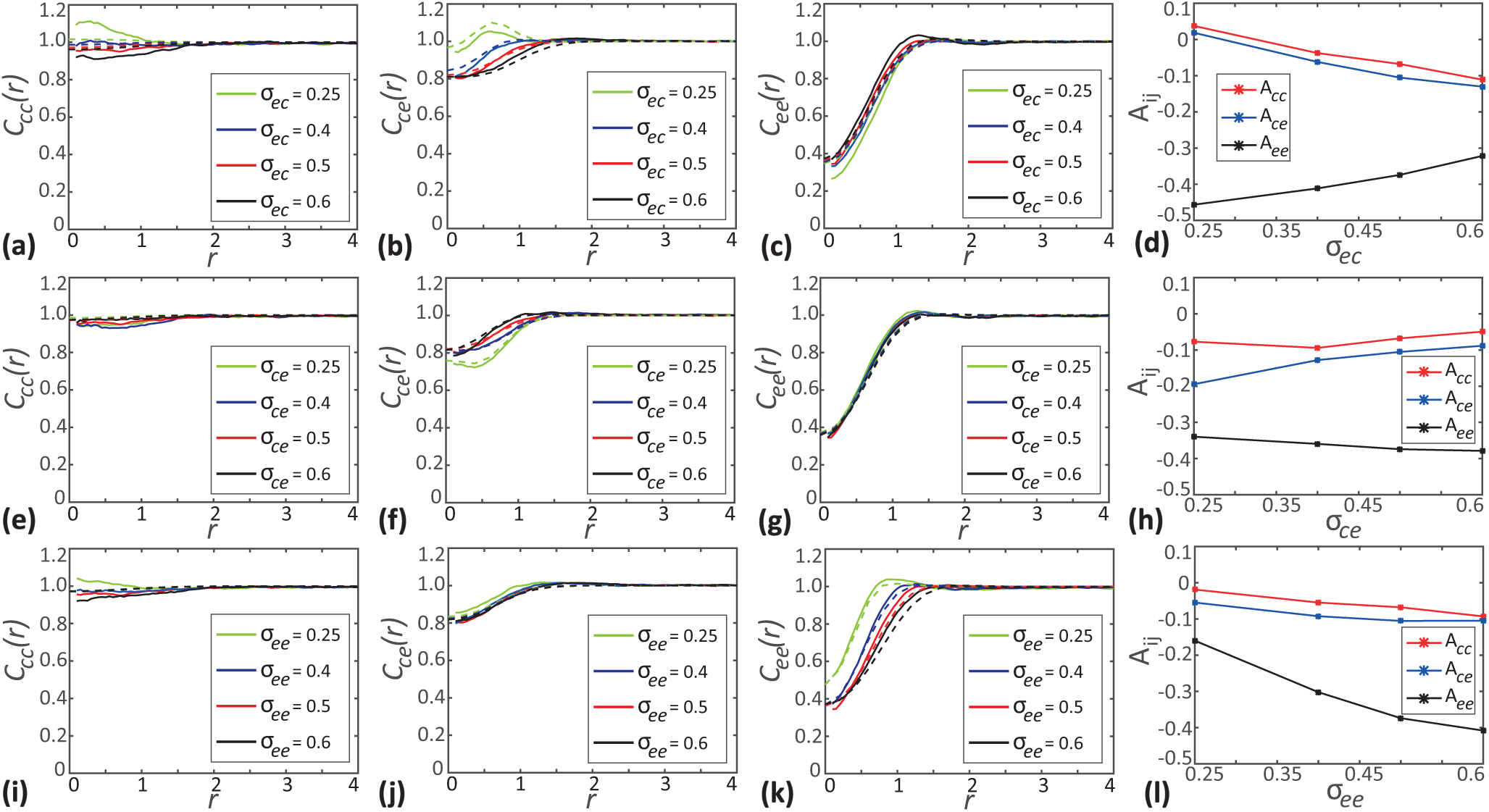
Effects of varying the spatial extent of interaction. **a-c** show the PCFs *C*_*cc*_(*r*), *C*_*ce*_(*r*) and *C*_*ee*_(*r*) as a function of separation distance for different choices of *σ*_*ec*_. **d** show simple measures of spatial structure, *A*_*cc*_, *A*_*ce*_ and *A*_*ee*_ as a function of *σ*_*ec*_. **e-g** shows *C*_*i j*_(*r*) for different choices of *σ*_*ce*_. **h** shows *A*_*i j*_ as a function of *σ*_*ce*_. **i-k** shows *C*_*i j*_(*r*) for different choices of *σ*_*ee*_. **l** shows *A*_*i j*_ as a function of *σ*_*ee*_. In the subfigures showing the PCFs as a function of *r*, solid curves show the averaged results from 1000 identically prepared realisations of the IBM, whereas dashed curves correspond to results from spatial moment dynamics model. In all cases *C*_*i j*_(*r*) and *A*_*i j*_ are evaluated at *t* = 20. Parameter values are *γ*_*cc*_ = 0, *γ*_*ce*_ = −0.2, *γ*_*ec*_ = *γ*_*ee*_ = 0.3, *m*_*c*_ = *m*_*e*_ = 5.0, 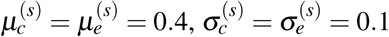.

In Figure 5(e)-(h) we vary *σ*_*ce*_, which controls the spatial extent over which chasers are attracted to escapees. At small values of *σ*_*ce*_ we observe an interspecies regular spatial structure between chasers and escapees as indicated by *C*_*ce*_(*r*) *<* 1 and *A*_*ce*_ *<* 0. For sufficiently small *σ*_*ce*_, the escapees are repelled from chasers strongly since the small range attraction of chasers to escapees cannot compensate for the relatively long range repulsion. Hence, chasers and escapees segregate from each other forming an interspecies regular spatial structure. As *σ*_*ce*_ increases, we observe the cross-PCF *C*_*ce*_(*r*) approaches unity and *A*_*ce*_ becomes less negative. This suggests a shift from the interspecies regular spatial structure between chasers and escapees to a less regular spatial structure. With the increase of the spatial extent of interaction, the distance over which chasers get attracted to escapees increases and results in an enhanced attractive bias. This enhanced attraction counteracts the interspecies repulsion leading to the loss of the regular spatial pattern between chasers and escapees.

Finally, in Figure 5(i)-(l) we vary *σ*_*ee*_, which controls the spatial extend over which escapees repel other escapees. As *σ*_*ee*_ increases, we see the *C*_*ee*_(*r*) diverges from unity and *A*_*ee*_ becomes more negative, suggesting the development of stronger regular spatial structure among escapees. The increase in *σ*_*ee*_ enables an escapee to influence more distant escapees, resulting in the enhancement of the regular structure.

### Effect of varying the relative density of species

We now explore how varying the relative density of chasers and escapees, *Z*_1,*c*_*/Z*_1,*e*_, impacts the spatial configuration of the community. We consider a community composed of three different density ratios: *Z*_1,*c*_*/Z*_1,*e*_ = 1*/*7, *Z*_1,*c*_*/Z*_1,*e*_ = 1 and *Z*_1,*c*_*/Z*_1,*e*_ = 1. To vary the relative density of chasers, the total number of individuals in the community is fixed at *N*_*c*_ + *N*_*e*_ = 400 and the population sizes of chasers are varied as *N*_1_ = 50, 200 and 350, respectively. We investigate the behaviour of spatial pattern in the community by plotting *C*_*i j*_(*r*) and *A*_*i j*_ in Figure 6.

**Figure 6.**
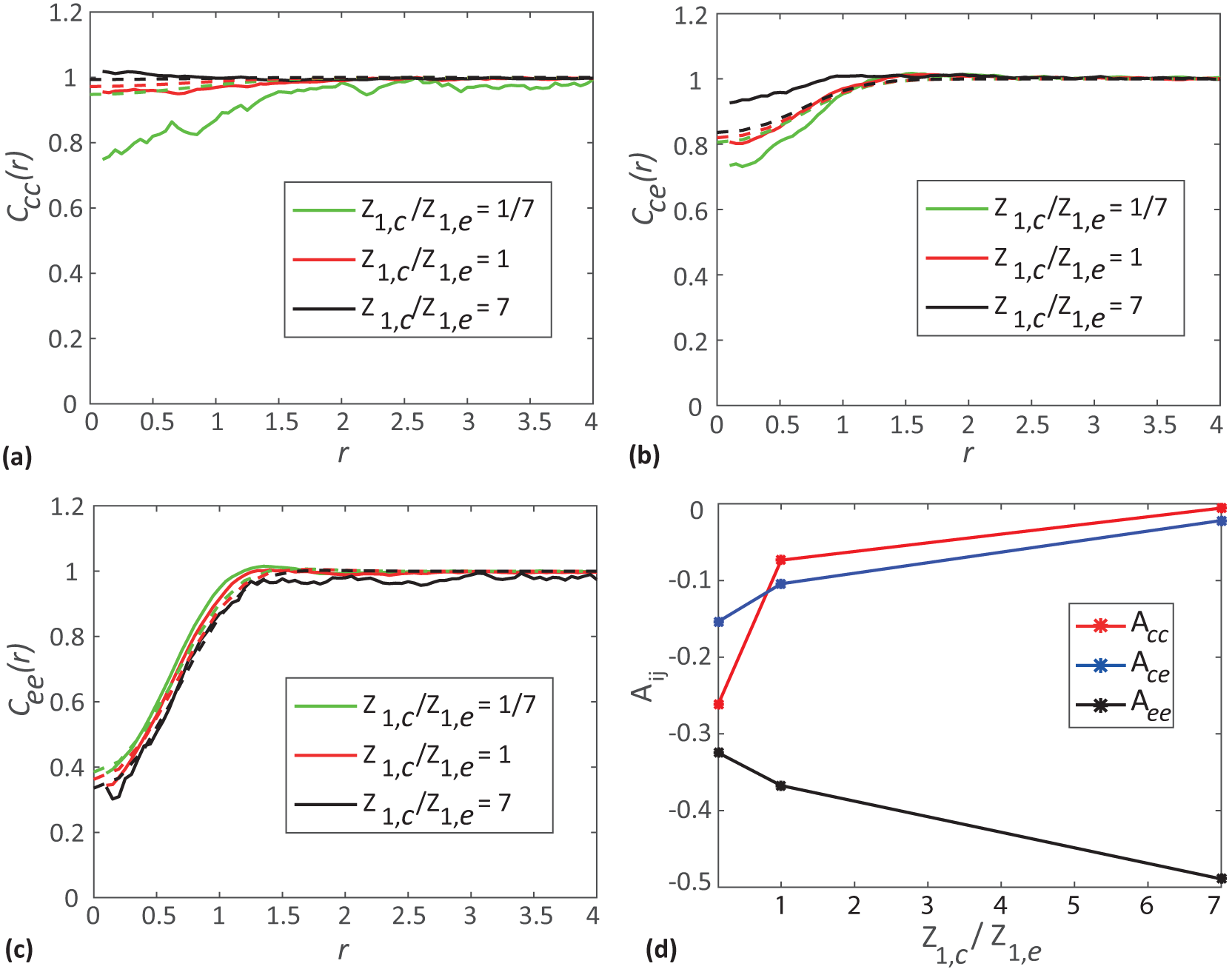
Effects of varying the relative density. **a-c** shows *C*_*cc*_(*r*), *C*_*ce*_(*r*) and *C*_*ee*_(*r*) as a function of separation distance, respectively. **d** show simple measures of spatial structure, *A*_*i j*_ as a function of relative density. In (**a**)-(**c**) solid curves show the averaged results from 1000 identically prepared realisations of the IBM, whereas dashed curves correspond to results from spatial moment dynamics model. In all results, PCFs and simple measures of spatial structure are given at *t* = 20. Parameter values are *γ*_*cc*_ = 0, *γ*_*ce*_ = −0.2, *γ*_*ec*_ = *γ*_*ee*_ = 0.3, *σ* = *σ* = *σ* = *σ* = 0.5, *m* = *m* = 5.0,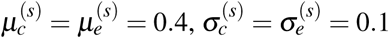.

When the density of chasers is much lower than the density of escapees, the attraction of chasers towards escapees leads to chasers being placed far apart from other chasers. This leads to the formation of regular spatial structure (*C*_*cc*_ *<* 1 and *A*_*cc*_ *<* 0) among chasers. In contrast, at a high relative density of chasers, more chasers tend to move towards a particular escapee, resulting in an accumulation of chasers at short distances and a less regular spatial structure, as indicated by *C*_*cc*_(*r*) becomes close to unity and *A*_*cc*_ becomes closer to zero. Another observation is that *C*_*ee*_(*r*) decreases and *A*_*ee*_ becomes more negative with the increase of relative density of chasers. This indicates the emergence of a stronger intraspecies regular spatial pattern among escapees. This is a potentially counterintuitive result since we see stronger regular structure in escapees as its population size decreases. This is an indirect effect of interactions of escapees with chasers. At a high relative density of chasers, each escapee will be surrounded by many chasers. The repulsive bias of escapees to chasers forbids the possible movement of any escapee, towards other escapees, that are surrounded by chasers. This results in the segregation of escapees. We see *C*_*ce*_(*r*) becomes close to one and *A*_*ce*_ approaches zero as we increase the relative density of chasers, suggesting the weakening of the interspecies regular spatial pattern. As we increase the relative density of chasers, the attraction of chasers to escapees outweighs the repulsive bias of escapees from chasers, to create a less regular spatial pattern. We also provide another set of results in Section 6 of the Supplementary Material to emphasise that the impact of variation in relative density is strongly dependent on the choice of interaction strengths.

## Conclusions

The most striking feature of our analysis is that we find the formation of, at times, a strong spatial structure that is driven purely by motility. This is interesting because most spatial moment models and analysis have their origin in plant ecology where birth and death, rather than motility, are the key mechanisms^23, 24, 26, 32^. More recent spatial moment models have been developed to mimic cell biology experiments where simultaneous movement and proliferation are important, and these previous studies attribute the formation of spatial structure to the combined effects of motility and proliferation^27, 28^. As far as we are aware, this is the first study to apply spatial moment analysis to a conservative community, composed of multiple distinct populations, without birth or death events. These considerations enable us to focus on how motility alone drives significant spatial patterning, and we find that this is particularly relevant when we consider asymmetric interactions between distinct populations within the community. Our IBM and spatial moment dynamics modelling framework allows us to capture different features of the individual-level motility mechanisms such as neighbour dependent directional bias emerging as a consequence of the interactions among individuals. We explore a range of complex spatial structure arising from the combined impacts of various intraspecies and interspecies interactions, which involve different types of attractive or repulsive bias between individuals in a stereotypical community consisting of two distinct species. We provide an extensive analysis of different features of the intraspecies and interspecies interaction by varying the interaction strength, the spatial extent of interaction and relative densities of individuals of both the species. Our results include several interesting and possibly counter-intuitive results. For example, under self-repulsion of escapees and chase-escape behaviour, the increase in density of chasers relative to escapees results in weakening of regular pattern between chasers and enhancement of regular pattern among escapees.

An interesting extension of the model presented here would be to include interactions which affect the proliferation and mortality of individuals, such as predator-prey interactions or intra and interspecific competition^44, 45^. The population dynamics resulting from such interactions will depend on the types of spatial structure examined in this work. Another extension would be to consider spatial processes such as moving fronts that are relevant to many biological processes including malignant invasion and developmental morphogenesis^46^. In this study, we consider the movement of individuals in a two-dimensional spatial domain. However, realistic modelling of *in vivo* cell migration requires consideration of three-dimensional (3D) space. While the extension of the IBM framework and moment dynamics analysis to 3D is relatively straight forward, the numerical evaluation of moment dynamics in 3D can become increasingly complicated. We leave these extensions for future work.

## Acknowledgements

This work is supported by the Australian Research Council (DP170100474). MJP is partly supported by Te Pūnaha Matatini, a New Zealand Centre of Research Excellence.

## Author contributions statement

AS, MJP, MJS conceived the study. AS carried out the stochastic simulations, performed mathematical derivations and wrote code to solve the governing equations. AS, MJP, MJS interpreted the results. AS wrote the manuscript. AS, MJP, MJS edited the manuscript and approved the final version of the manuscript.

