## Supplementary Material for "Spatial structure arising from chase-escape interactions with crowding"

#### Contents

**1 Derivation of Movement displacement PDF 2**

**2 Spatial moment dynamics for a community consisting of two distinct species 3**

**3 Moment closure schemes 5**

**4 Evolution of spatial structure 7**

**5 Dependence of steady-state spatial patterns on the initial configuration of individuals 9**

**6 Effect of varying the relative density of species 11**

### 1 Derivation of Movement displacement PDF

Here, we derive an expression for the probability density function (PDF), for the displacement that a chaser traverses during a movement event. Let us assume that the distance moved is a random variable with PDF $u_c(|\boldsymbol{\xi}|)$ and the direction of movement is specified by an independent random variable with PDF $g(\theta)$. Now, we have the movement vector, $(\boldsymbol{\xi}_x, \boldsymbol{\xi}_y) = (|\boldsymbol{\xi}|\cos(\theta), |\boldsymbol{\xi}|\sin(\theta))$ and the PDF for this bivariate random variable can be computed as,

$$\begin{aligned}
\mu_c(\boldsymbol{\xi}) &= \frac{\mathbb{P}(\text{movement vector}\,(\boldsymbol{\xi}_x, \boldsymbol{\xi}_y)\,\text{is in a region of area}\,\mathrm{dA} \in \mathbb{R}^2)}{\mathrm{dA}}, \\
&= \frac{u_c(|\boldsymbol{\xi}|)\mathrm{d}|\boldsymbol{\xi}| \times \mathrm{g}(\theta)\mathrm{d}\theta}{|\boldsymbol{\xi}|\,\mathrm{d}|\boldsymbol{\xi}|\,\mathrm{d}\theta}, \\
&= \frac{u_c(|\boldsymbol{\xi}|)\,g(\theta)}{|\boldsymbol{\xi}|}.
\end{aligned} \tag{S1}$$

A similar derivation holds for the movement displacement PDF, $\mu_e(\boldsymbol{\xi})$, of escapees.

#### 2 Spatial moment dynamics for a community consisting of two distinct species

Now, we present the details about the spatial moment dynamics for a specific case of the generalised model, where the community consists of individuals from two distinct species. The average densities of chasers and escapees are denoted by $Z_{1,c}$ and $Z_{1,e}$, respectively. Four different second spatial moments correspond to the average density of pairs of individuals are denoted by $Z_{2,cc}(\boldsymbol{\xi}, t)$, $Z_{2,ce}(\boldsymbol{\xi}, t)$, $Z_{2,ec}(\boldsymbol{\xi}, t)$, and $Z_{2,ee}(\boldsymbol{\xi}, t)$, respectively.

The gradient of the bias kernel gives the contribution of individuals of both species to the bias vector of the neighbouring individual. The expected net bias vector of a chaser conditional on the presence of another chaser is given by,

$$\begin{aligned}\mathbf{B}_{2,cc}(\boldsymbol{\xi}, t) = \frac{1}{Z_{2,cc}(\boldsymbol{\xi}, t)} \int \Big( &\nabla\omega_{cc}(|\boldsymbol{\xi}'|) Z_{3,ccc}(\boldsymbol{\xi}, \boldsymbol{\xi}', t) \\ &+ \nabla\omega_{ce}(|\boldsymbol{\xi}'|) Z_{3,cce}(\boldsymbol{\xi}, \boldsymbol{\xi}', t)\Big)\, \mathrm{d}\boldsymbol{\xi}' + \nabla\omega_{cc}(|\boldsymbol{\xi}|).\end{aligned} \tag{S2}$$

Similarly, the expected net bias vector of a chaser conditional on the presence of an escapee is given by,

$$\begin{aligned}\mathbf{B}_{2,ce}(\boldsymbol{\xi}, t) = \frac{1}{Z_{2,ce}(\boldsymbol{\xi}, t)} \int \Big( &\nabla\omega_{cc}(|\boldsymbol{\xi}'|) Z_{3,cec}(\boldsymbol{\xi}, \boldsymbol{\xi}', t) \\ &+ \nabla\omega_{ce}(|\boldsymbol{\xi}'|) Z_{3,cee}(\boldsymbol{\xi}, \boldsymbol{\xi}', t)\Big)\, \mathrm{d}\boldsymbol{\xi}' + \nabla\omega_{ce}(|\boldsymbol{\xi}|).\end{aligned} \tag{S3}$$

The expected net bias vectors of an escapee conditional on the presence of chaser and escapee, respectively are given by,

$$\begin{aligned}\mathbf{B}_{2,ec}(\boldsymbol{\xi}, t) = \frac{1}{Z_{2,ec}(\boldsymbol{\xi}, t)} \int \Big( &\nabla\omega_{ec}(|\boldsymbol{\xi}'|) Z_{3,ecc}(\boldsymbol{\xi}, \boldsymbol{\xi}', t) \\ &+ \nabla\omega_{ee}(|\boldsymbol{\xi}'|) Z_{3,ece}(\boldsymbol{\xi}, \boldsymbol{\xi}', t)\Big)\, \mathrm{d}\boldsymbol{\xi}' + \nabla\omega_{ec}(|\boldsymbol{\xi}|),\end{aligned} \tag{S4}$$

$$\begin{aligned}\mathbf{B}_{2,ee}(\boldsymbol{\xi}, t) = \frac{1}{Z_{2,ee}(\boldsymbol{\xi}, t)} \int \Big( &\nabla\omega_{ec}(|\boldsymbol{\xi}'|) Z_{3,eec}(\boldsymbol{\xi}, \boldsymbol{\xi}', t) \\ &+ \nabla\omega_{ee}(|\boldsymbol{\xi}'|) Z_{3,eee}(\boldsymbol{\xi}, \boldsymbol{\xi}', t)\Big)\, \mathrm{d}\boldsymbol{\xi}' + \nabla\omega_{ee}(|\boldsymbol{\xi}|).\end{aligned} \tag{S5}$$

We now develop the equations governing the dynamics of the second moments. The equations for the density of pairs depends on the loss and gain of pairs at displacement $\boldsymbol{\xi}$. The equations governing the dynamics of second moments for the community of chasers and escapees are given by,

$$\begin{aligned}\frac{\partial}{\partial t} Z_{2,cc}(\boldsymbol{\xi}, t) = & - 2\, m_c\, Z_{2,cc}(\boldsymbol{\xi}, t) \\ & + m_c \int \mu_{2,cc}(\boldsymbol{\xi'}, \boldsymbol{\xi'} + \boldsymbol{\xi}, t)\, Z_{2,cc}(\boldsymbol{\xi'} + \boldsymbol{\xi}, t)\, \mathrm{d}\boldsymbol{\xi'} \\ & + m_c \int \mu_{2,cc}(\boldsymbol{\xi'}, \boldsymbol{\xi'} - \boldsymbol{\xi}, t)\, Z_{2,cc}(\boldsymbol{\xi'} - \boldsymbol{\xi}, t)\, \mathrm{d}\boldsymbol{\xi'},\end{aligned} \tag{S6}$$

$$\begin{aligned}\frac{\partial}{\partial t} Z_{2,ce}(\boldsymbol{\xi}, t) = & - \big(m_c + m_e\big) Z_{2,ce}(\boldsymbol{\xi}, t) \\ & + m_c \int \mu_{2,ce}(\boldsymbol{\xi'}, \boldsymbol{\xi'} + \boldsymbol{\xi}, t)\, Z_{2,ce}(\boldsymbol{\xi'} + \boldsymbol{\xi}, t)\, \mathrm{d}\boldsymbol{\xi'} \\ & + m_e \int \mu_{2,ec}(\boldsymbol{\xi'}, \boldsymbol{\xi'} - \boldsymbol{\xi}, t)\, Z_{2,ec}(\boldsymbol{\xi'} - \boldsymbol{\xi}, t)\, \mathrm{d}\boldsymbol{\xi'},\end{aligned} \tag{S7}$$

$$\begin{aligned}\frac{\partial}{\partial t} Z_{2,ec}(\boldsymbol{\xi}, t) = & - \big(m_c + m_e\big) Z_{2,ec}(\boldsymbol{\xi}, t) \\ & + m_e \int \mu_{2,ec}(\boldsymbol{\xi'}, \boldsymbol{\xi'} + \boldsymbol{\xi}, t)\, Z_{2,ec}(\boldsymbol{\xi'} + \boldsymbol{\xi}, t)\, \mathrm{d}\boldsymbol{\xi'} \\ & + m_c \int \mu_{2,ce}(\boldsymbol{\xi'}, \boldsymbol{\xi'} - \boldsymbol{\xi}, t)\, Z_{2,ce}(\boldsymbol{\xi'} - \boldsymbol{\xi}, t)\, \mathrm{d}\boldsymbol{\xi'},\end{aligned} \tag{S8}$$

$$\begin{aligned}\frac{\partial}{\partial t} Z_{2,ee}(\boldsymbol{\xi}, t) = & - 2\, m_e\, Z_{2,ee}(\boldsymbol{\xi}, t) \\ & + m_e \int \mu_{2,ee}(\boldsymbol{\xi'}, \boldsymbol{\xi'} + \boldsymbol{\xi}, t)\, Z_{2,ee}(\boldsymbol{\xi'} + \boldsymbol{\xi}, t)\, \mathrm{d}\boldsymbol{\xi'} \\ & + m_e \int \mu_{2,ee}(\boldsymbol{\xi'}, \boldsymbol{\xi'} - \boldsymbol{\xi}, t)\, Z_{2,ee}(\boldsymbol{\xi'} - \boldsymbol{\xi}, t)\, \mathrm{d}\boldsymbol{\xi'}.\end{aligned} \tag{S9}$$

#### 3 Moment closure schemes

Here, we compare the accuracy of four popular moment closure schemes. The closure schemes considered include the power-1 closure (P1), symmetric power-2 closure (P2S), asymmetric power-2 closure (P2A) and Kirkwood superposition approximation (KSA) [1, 2].

The power-1 closure (P1) is given by,

$$\begin{aligned} Z_{3,ijk}(\boldsymbol{\xi}, \boldsymbol{\xi}', t) =& Z_{1,i}\, Z_{2,jk}(\boldsymbol{\xi}' - \boldsymbol{\xi}, t) + Z_{1,j}\, Z_{2,ik}(\boldsymbol{\xi}', t) \\ & + Z_{1,k}\, Z_{2,ij}(\boldsymbol{\xi}, t) - 2 Z_{1,i}\, Z_{1,j}\, Z_{1,k}. \end{aligned} \tag{S10}$$

The power-2 closure is given by,

$$\begin{aligned} Z_{3,ijk}(\boldsymbol{\xi}, \boldsymbol{\xi}', t) =& \frac{1}{\alpha + \gamma} \Big[ \alpha \frac{Z_{2,ij}(\boldsymbol{\xi}, t)\, Z_{2,ik}(\boldsymbol{\xi}', t)}{Z_{1,i}} + \beta \frac{Z_{2,ij}(\boldsymbol{\xi}, t)\, Z_{2,jk}(\boldsymbol{\xi}' - \boldsymbol{\xi}, t)}{Z_{1,j}} \\ & + \gamma \frac{Z_{2,ik}(\boldsymbol{\xi}', t)\, Z_{2,jk}(\boldsymbol{\xi}' - \boldsymbol{\xi}, t)}{Z_{1,k}} - \beta Z_{1,i} Z_{1,j} Z_{1,k} \Big]. \end{aligned} \tag{S11}$$

For the symmetric power-2 closure (P2S), we choose $\alpha = \beta = \gamma = 1$, and for the asymmetric power-2 closure (P2A), we use $\alpha = 4, \beta = 1$, and $\gamma = 1$ [3].

The Kirkwood superposition approximation (KSA) is given by,

$$Z_{3,ijk}(\boldsymbol{\xi}, \boldsymbol{\xi}', t) = \frac{Z_{2,ij}(\boldsymbol{\xi}, t)\, Z_{2,ik}(\boldsymbol{\xi}', t)\, Z_{2,jk}(\boldsymbol{\xi}' - \boldsymbol{\xi}, t)}{Z_{1,i}\, Z_{1,j}\, Z_{1,k}}. \tag{S12}$$

We compute PCFs using these different closure schemes for a community consisting of two species with $N_c = N_e = 100$ and compare with the averaged results of IBM simulation. Again, in these simulations, we consider a randomly distributed initial arrangement of individuals. Results in Figure S1 compare the auto-PCFs, $C_{cc}(r)$ and $C_{ee}(r)$, and the cross-PCF, $C_{ce}(r)$, for each of the closure schemes with results from the IBM simulation. Overall, when we consider all three PFCs, we see that the power-2 asymmetric closure provides the best match with the prediction of the IBM simulation for the parameters considered in Figure S1. Our approach in this work is not to promote one particular closure approximation over another closure approximation. Instead, we present a general spatial moment framework that can be implemented with a range of closure assumptions should one particular closure approximation be preferred to another.

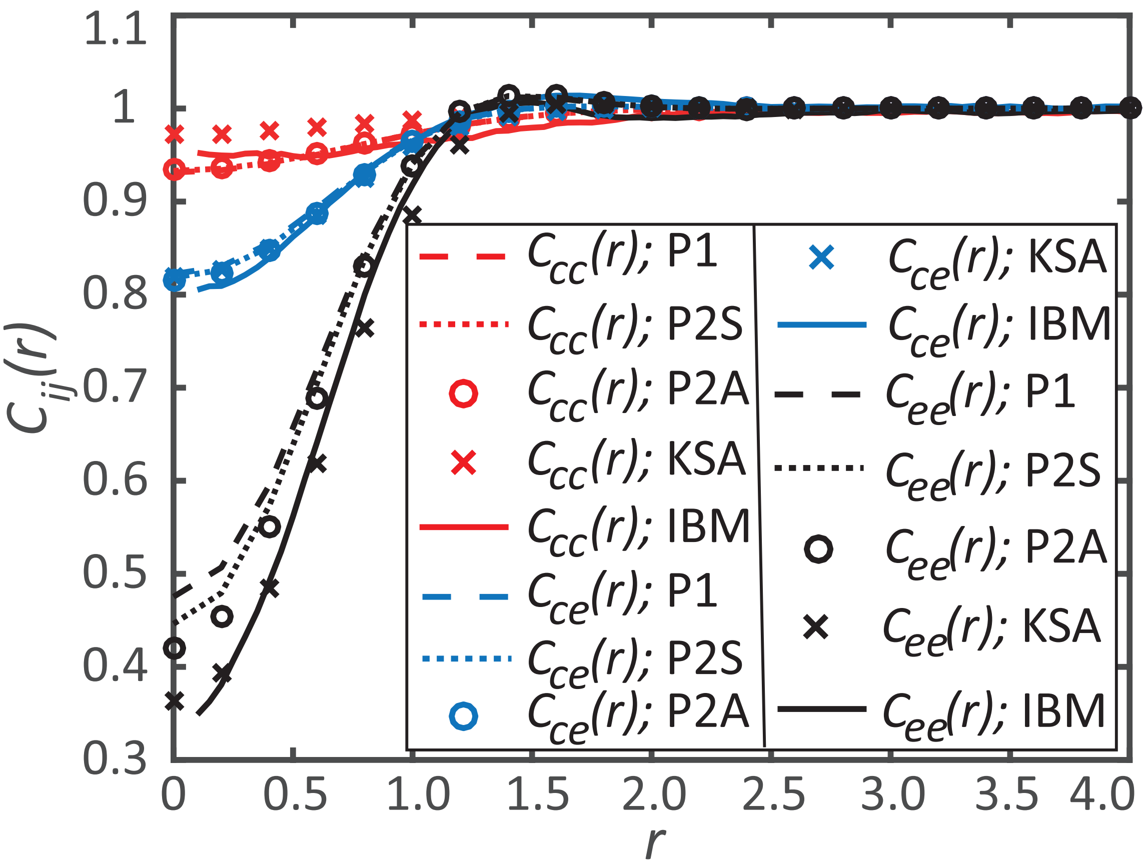


Figure S1: Comparison of spatial moment closure approximation schemes. The curves in red, blue and black correspond to $C_{cc}(r), C_{ce}(r)$ and $C_{ee}(r)$, respectively. In all results, PCFs are given at $t = 20$. All PCFs from IBM correspond to averaged data of 1000 identically prepared realisations. Parameter values are $\gamma_{cc} = 0, \gamma_{ce} = -0.2, \gamma_{ec} = \gamma_{ee} = 0.3$, $\sigma_{cc} = \sigma_{ce} = \sigma_{ec} = \sigma_{ee} = 0.5$, $m_c = m_e = 5.0$, $\mu_c^{(s)} = \mu_e^{(s)} = 0.4$, $\sigma_c^{(s)} = \sigma_e^{(s)} = 0.1$.

### 4 Evolution of spatial structure

Here, we explore how the spatial structure evolve as individuals undergo movement events. To illustrate, we consider a sample community consisting of individuals from two distinct species with population sizes $N_c = 30$ and $N_e = 30$. In this suite of simulations, we use a negative $\gamma_{ce}$, meaning that the chasers are attracted to escapees. All remaining bias strength parameters are chosen to be positive, which correspond to conspecific repulsions and repulsion of escapees to chasers. Results in Figure S2 show a series of snapshots of the location of individuals and PCFs computed at $t = 0, 1, 20$, and 40, respectively.

As time progresses, the locations of individuals change due to the movement of individuals as shown in Figure S2(a)-(d). That implies, the local environment around individuals keeps changing and hence the neighbourhood contribution to the bias vector of an individual also varies. The transition of the spatial structure of the community from the initial spatial Poisson process is explored by plotting the PCFs $C_{cc}(r), C_{ee}(r)$ and $C_{ce}(r)$, respectively as shown in Figure S2(e)-(h). Comparing the results at $t = 20$ and $t = 40$, suggests that we have an approximately steady spatial pattern established by $t = 20$. As time progresses, we observe the development of a regular spatial pattern among subpopulations of chasers and escapees since $C_{cc}(r) < 1$ and $C_{ee}(r) < 1$ for small distances. The reason for the regular spatial structure is the intraspecies repulsion among both species. Each chaser (escapee) tries to stay apart from other chasers (escapees), hence leading to a regular spatial structure among chasers (escapees). We also observe a less pronounced interspecies regular spatial pattern between chasers and escapees. Here, the attractive bias of chasers to escapees counteracts the repulsive bias of escapees to chasers, reducing the extent of the interspecies regular spatial pattern.

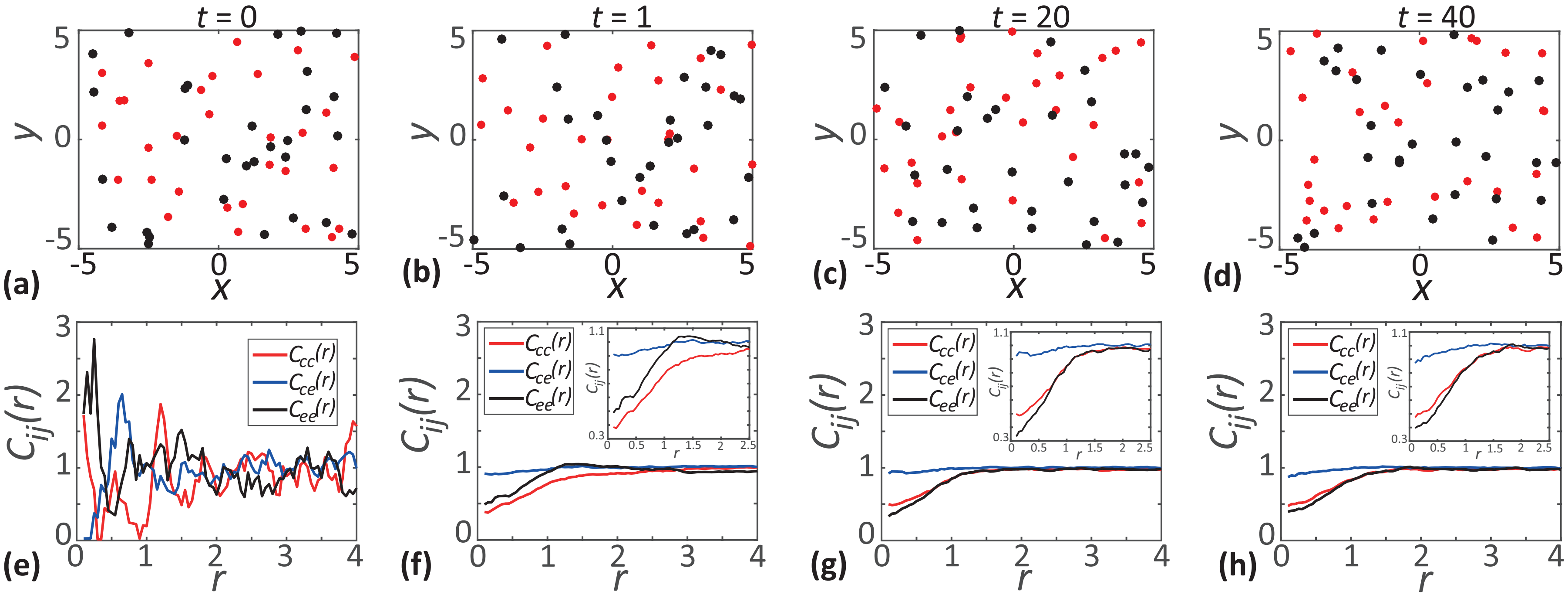


Figure S2: Results in **a-d** show the location of chasers (red dots) and escapees (black dots) at $t = 0, 1, 20$ and 40, respectively. Results in **e-h** show the PCFs $C_{cc}(r)$, $C_{ce}(r)$, $C_{ee}(r)$ as a function of separation distance $r$ at $t = 0, 1, 20$ and 40, respectively. The PCFs in **e** corresponds to the randomly distributed initial arrangement of individuals and all PCFs in **f-h** are computed using the average data from 1000 identically prepared realisations of the IBM. Parameter values are $\gamma_{cc} = 0.1, \gamma_{ce} = -0.1$, $\gamma_{ec} = \gamma_{ee} = 0.15$ and $\sigma_{cc} = \sigma_{ce} = \sigma_{ec} = \sigma_{ee} = 0.5$.

### 5 Dependence of steady-state spatial patterns on the initial configuration of individuals

In this section we present additional results which show that the steady-state spatial structure is independent of the initial distribution of individuals. To illustrate, we consider two different initial configurations shown in Figure S3(a)-(b), respectively. Here, the first case corresponds to a uniform distribution of chasers and escapees with population sizes, $N_c = 200$ and $N_e = 200$, respectively. The second case considered corresponds to a spatially non-uniform distribution of chasers and escapees. In this second case, chasers and escapees are placed at the opposite corners of the domain. The population sizes of both the species are kept the same as that in Figure S3(a). To examine whether these different initial conditions affect the long-time spatial structure we simulate the IBM with these two different types of initial distributions of individuals and in both types of simulations we use identical interaction strengths and spatial extents of interactions.

We observe that a steady spatial pattern emerges in the case of simulation with uniform initial arrangement of chasers and escapees by approximately $t = 20$, whereas the second initial condition requires a longer duration to approach a steady distribution, $t = 100$. To examine the resulting spatial structure we plot the auto and cross-PCFs in Figure S3(c) and these results confirm that the two different initial configurations leads to identical spatial structure.

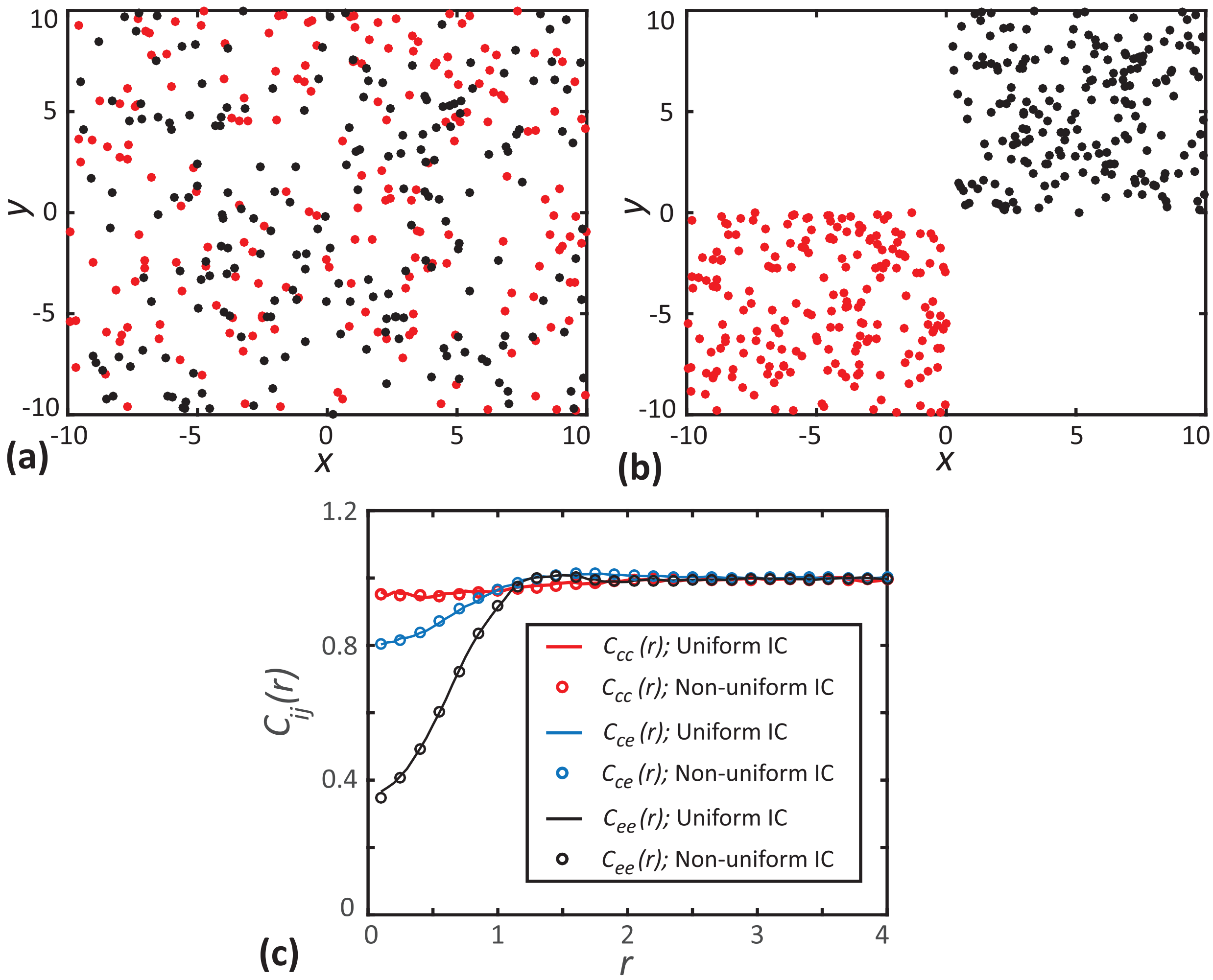


Figure S3: Effect of varying the initial configuration of individuals on the spatial structure. **a** shows a uniformly distributed initial arrangement of chasers (red dots) and escapees (black dots). **b** shows a non-uniform initial arrangement of individuals. In this case, chasers and escapees are placed at the opposite corners of the domain. **c** shows the auto and cross-PCFs computed with uniform and non-uniform initial configurations (IC) at $t = 20$ (solid lines) and $t = 100$ (open circles), respectively. In **c** red, blue and black curves represent $C_{cc}(r)$, $C_{ce}(r)$ and $C_{ee}(r)$, respectively. Parameter values are $\gamma_{cc} = 0, \gamma_{ce} = -0.2$, $\gamma_{ec} = \gamma_{ee} = 0.3$ and $\sigma_{cc} = \sigma_{ce} = \sigma_{ec} = \sigma_{ee} = 0.5$.

### 6 Effect of varying the relative density of species

Here, we present another set of results, as shown in Figure S4, to emphasise that the impact of variation in relative density is strongly dependent on the choice of interaction strengths. To illustrate, we consider a scenario with same parameters and initial placement of individuals as in Figure 7 in the Main Paper, except that we use a positive $\gamma_{ce}$ that corresponds to repulsion of chasers from escapees. In contrast to the results in Figure 7 in the Main Paper, we observe a strong clustering of chasers at low relative densities of chasers. The presence of more escapees creates a strong repulsion to fewer chasers and the choice of $\gamma_{ce} = 0.3$ forces chasers to move away from the escapees. Since there is no interaction between chasers, both these factors act together to form clusters of chasers. As we increase the relative density, the clustering weakens and leads to a random spatial structure. This is also confirmed by the lowering of simple measure of spatial structure, $A_{cc}$, from 1.5 at $Z_{1,c}/Z_{1,e} = 1/7$ to close to zero at $Z_{1,c}/Z_{1,e} = 7$.

At low relative densities of chasers we observe an intraspecies regular spatial pattern of escapees due to the repulsive interaction between escapees. As we increase the relative density of chasers, we observe a peak in PCF, $C_{ee}(r)$, at approximately $r = 1.25$. This peak corresponds to an extreme case of regular spatial pattern where almost all escapees are separated by approximately the same distance. The more pronounced regular pattern arises due to the combined effect of the enhanced repulsion of chasers to escapees, due to the increase in the number of chasers compared to escapees, and the repulsive intraspecies interaction of escapees. Overall, we summarise that the interspecies regular spatial structure between chasers and escapees becomes more pronounced as the relative density of chasers increases.

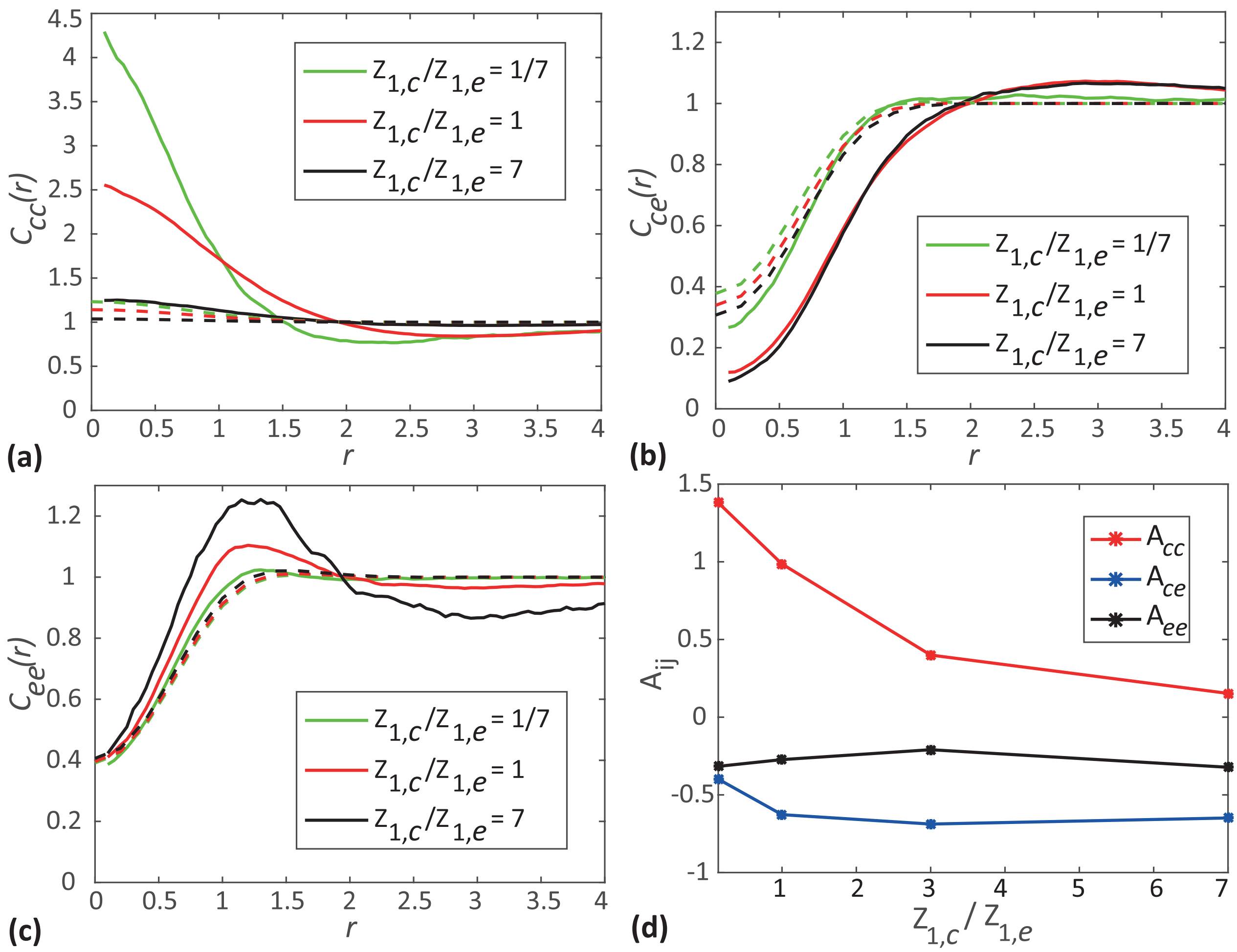


Figure S4: Effects of varying the relative density of species. **a** shows the auto-PCF of chasers as a function of separation distance. **b** shows the cross-PCF as a function of separation distance. **c** shows the auto-PCF of escapees as a function of separation distance. **d** show simple measures of spatial structure, $A_{ij}$, as a function of relative densities of species. In (**a**)-(**c**) solid curves show the averaged results from 1000 identically prepared realisations of the IBM, whereas dashed curves correspond to results from spatial moment dynamics model. In all results, PCFs and simple measures of spatial structure are given at $t = 20$. Parameter values are $\gamma_{cc} = 0, \gamma_{ce} = \gamma_{ec} = \gamma_{ee} = 0.3$, $\sigma_{cc} = \sigma_{ce} = \sigma_{ec} = \sigma_{ee} = 0.5$, $m_c = m_e = 5.0$, $\mu_c^{(s)} = \mu_e^{(s)} = 0.4$, $\sigma_c^{(s)} = \sigma_e^{(s)} = 0.1$.
